# Duplications and functional specialization force distinct evolution of isoflavonoid biosynthetic genes in legumes

**DOI:** 10.1101/361386

**Authors:** Yin Shan Jiao, Yu Zhao, Wen Feng Chen

## Abstract

Isoflavonoids are specialized plant metabolites, almost exclusive to legumes, and synthesized by the phenylpropanoid pathway. Leguminous plants produce 5-deoxyflavonoids and 5-deoxyisoflavonoids that act in symbiosis with nitrogen-fixing bacteria and involved in plant pathogen and stress response. However, little is known about evolutional origin of legume-specific isoflavonoid biosynthesis pathway. Here, we explored the genome-wide analysis of key genes: chalcone synthase (CHS), chalcone reductase (CHR), isoflavone synthase (IFS) and isoflavone reductase (IFR), encoding enzymes involved in the biosynthesis of (iso) flavonoids in legumes and nonlegumes. Among them, *CHS*, *CHR* and *IFR* comprise multigene families, underling the significant role of gene duplication in the evolutionary. Most duplications of *CHS* were highly the conventional leguminous type, whereas some were grouped with nonleguminous *CHS* genes. We also found that *CHR* homologs in soybean and *Sesbania rostrata* previously reported were ambiguous and should be re-identified. Phylogenetic analysis and protein sequences alignment indicated that IFSs in legumes are highly conserved. Intriguingly, unlike other IFRs in legumes, IFR-like homologs in *Sophora flavescens* and *Lupinus* angustifolius shared high sequence similarity and protein structures with homologs in nonlegumes. Overall, these results offer reasonable gene annotations and comparative analysis and also provided a glimpse into evolutional route of legume-specific isoflavonoid biosynthesis.

**Highlight:** Isoflavonoids are specialized plant metabolites, almost exclusive to legumes. We firstly provide evidence that evolutional origin of legume-specific isoflavonoid biosynthesis may be driven by gene duplications and functional specialization.

## Introduction

Flavonoids including flavones, flavanols and condensed tannins are plant secondary metabolites and ubiquitous in nature. More than 9,000 different flavonoids have been identified in plants, and a particular subset of them is involved in mediating host specificity in legumes (Ferrer *et al*., 2008; Perret *et al*., 2000). Isoflavonoids, a class of flavonoids, form a group of distinct secondary metabolites distributed almost exclusively in legume family. Many legumes produce specific flavonoids that only induce Nod factor production in homologous rhizobia, and therefore act as important determinants of host range (Liu and Murray, 2016). Daidzein and genistein, two isoflavonoids produced by soybean (*Glycine max*), are crucial signaling molecules that induce *Bradyrhizobium japonicum* to secret Nod factors to establish the symbiosis. *Medicago truncatula*, a model legume that has symbiosis-specific relationship with rhizobia, is only nodulated by either *Sinorhizobium meliloti* or *Sinorhizobium medicae* (Andrews and Andrews, 2017). On the contrary, more than 30 rhizobial species can establish symbiosis with *Sophora flavescens*, the most broad-spectrum host (Jiao *et al*., 2015). Several studies have shown that the production and release of flavonoids is central to how host-symbiont specificity is achieved (Den Herder *et al*., 2007; Perret *et al*., 2000). However, knowledge about genetic gene resources involved in the biosynthetic pathways of flavonoids in legume is limited and evolutional comparative analysis are lacking.

Vestitol, an isoflavan phytoalexin produced by *Lotus japonicus*, is postulated to be synthesized as part of the phenylpropanoid pathway (Fig. 1). It has multiple branches common to non-legumes as well as legumes, which provide general flavonoids such as condensed tannins and isoflavonoids including glyceollins, medicarpin and vestitol. In *L*. *japonicus*, the cDNAs encoding 10 enzymes involved in the biosynthesis of vestitol have been identified, except 7,20-dihydroxy-40-O-methoxyisoflavanol dehydratase. Chalcone synthase (CHS) is a key enzyme that catalyzes the first committed step towards flavonoid biosynthesis. Base on phylogenetic analysis, legumes have had a further expansion of one branch of the CHS gene family (Zavala and Opazo, 2015). Isoliquiritigenin is produced by a legume-specific enzyme, chalcone reductase (CHR), acts on an intermediate of the CHS reaction, yielding deoxychalcone from the coupled catalytic action with CHS (Wang, 2011). CHR belongs to the aldo-keto reductase (AKR) superfamily (Jez *et al*., 1997). All members of this superfamily contain a catalytic tetrad of Asp-53, Tyr-58, Lys-87, and His-120 and a common NAD(P)H binding site that is located in a deep, large and hydrophobic pocket at the C-terminus end (Bomati *et al*., 2005). Recently, Caroline J. Sepiol *et al*. performed a genome-wide search of AKR gene family members and identified 14 CHR genes in soybean (Sepiol *et al*., 2017). Up to now, CHR enzymes have been identified in various leguminous plant species, including *Medicago sativa* (Ballance and Dixon, 1995), *Sesbania rostrata* (Goormachtig *et al*., 1999), *Pueraria montana* var. *lobata* (He *et al*., 2011), *Glycyrrhiza glabra* (Hayashi *et al*., 2013), and *L*. *japonicus* (Shimada *et al*., 2007a).

**Fig.1.**
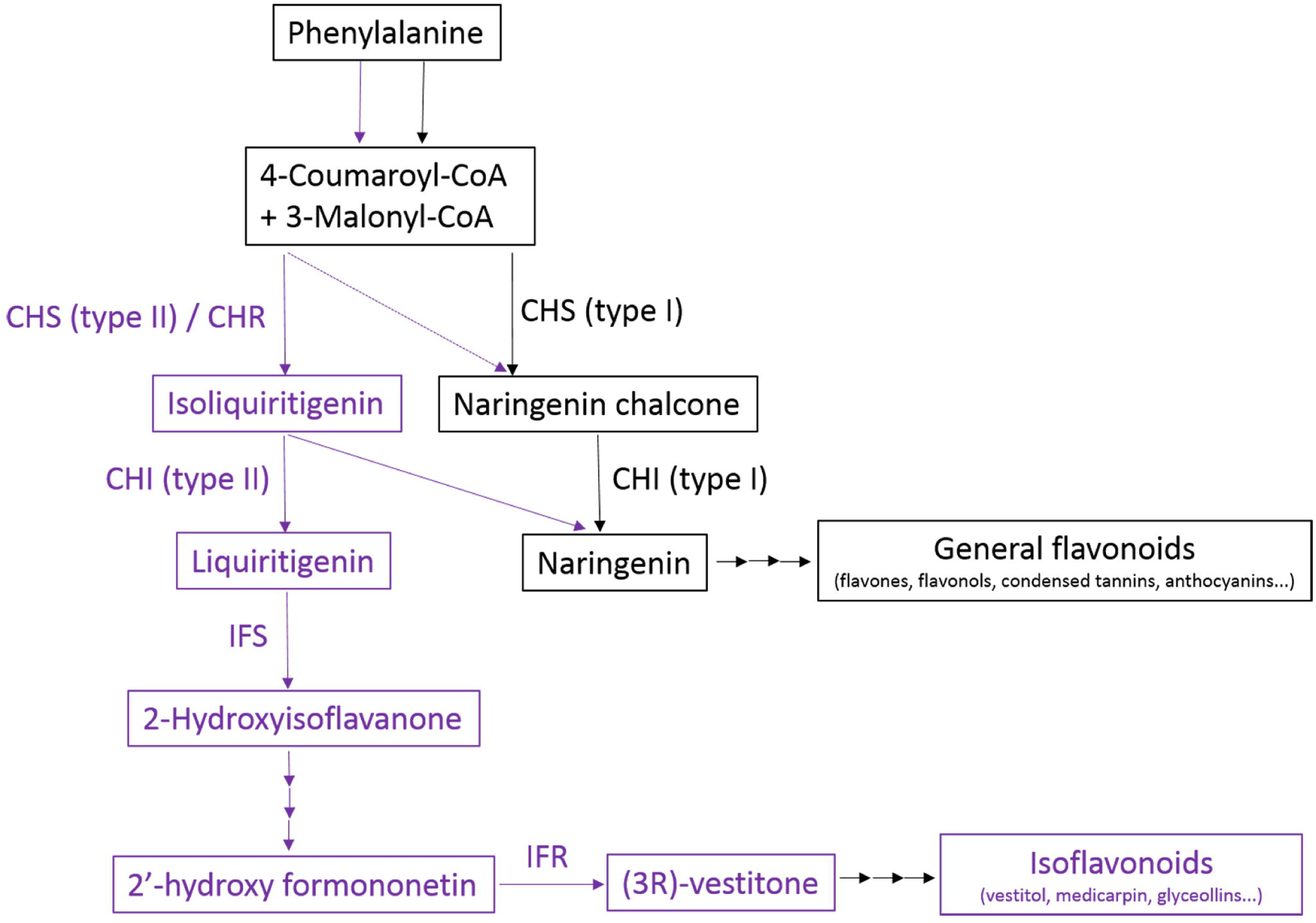
General and legume-specific (iso)flavonoid pathway in leguminous plants. Similar to CHIs, CHSs also have two types, legume-specific type (type II) and nonlegume type (type I). The schematic of biosynthetic pathway leading to isoflvonoids is shown in pink. The dotted arrow indicates speculative steps while multiple arrows indicate two or more steps in the pathway. CHR, chalcone reductase; CHS, chalcone synthase; CHI, chalcone isomerase; IFS, isoflavone synthase.

Isoflavone synthase (IFS), a key cytochrome P450 enzyme, is the first step in the branch of the phenylpropanoid pathway that commits metabolic intermediates to the synthesis of isoflavones (Wang, 2011). *Arabidopsis* possesses 244 putative P450 genes (and 28 pseudogenes), but none of them has IFS activity for synthesizing isoflavone (Bak *et al*., 2011). In soybean, IFS1 and IFS2 were identified by using a soybean EST collection (Jung *et al*., 2000). Introducing soybean IFS1 into *Arabidopsis thaliana* resulted in production of genistein (Jung *et al*., 2000), indicating its central role in the biosynthesis of isoflavonoids and extremely high substrate specificity. Interestingly, a recent study has shown that both IFS1 and IFS2 show positive interaction with upstream enzymes involved in isoflavonoid biosynthesis in soybean (Dastmalchi *et al*., 2016). Isoflavone reductase (IFR) was first identified as a key enzyme involved in the latter part of the medicarpin biosynthetic pathway (Guo *et al*., 1994; Paiva *et al*., 1991). IFR converts 2’-hydroxyformononetin to (3R)-vestitone (Fig. 1). In *M*. *truncatula*, MtIFR was first identified as a key enzyme involved in the biosynthetic pathway of the medicarpin, a major isoflavonoid phytoalexin (Paiva et al., 1991). Similar function of IFR homologs were also reported in chickpea (*Cicer arietinum*) and pea (*Pisum sativum*) (Daniel *et al*., 1990; Sun *et al*., 1991). Recently, a novel member of the soybean isoflavone reductase gene family *GmIFR* was identified (Cheng *et al*., 2015). Over expression of *GmIFR* transgenic soybean exhibited enhanced resistance to *Phytophthora*. Although isoflavonoid phytoalexins are restricted primarily to legumes, IFR-like proteins have been isolated from various nonlegumes such as maize (*Zea mays*) (Petrucco *et al*., 1996), rice (*Oryza saliva*) (Kim *et al*., 2003) and tobacco (*Nicotiana tabacum*) (Shoji *et al*., 2002). Here, we screened the available genomic and transcriptional resources of legumes distributed in three phylogenetic lineages to identify candidate genes encoding the four key enzymes: CHS, CHR, IFS and IFR. Phylogenetic relationship, comparative analysis of nucleotide and protein sequences and evolutional processes of these four gene homologs in legumes and nonlegumes, were investigated. We demonstrate that: (a) similar to *CHI*, *CHS* also has legume-specific type and evolved more rounds of gene duplications compared to those in nonlegumes; (b) *CHR* homologs in soybean and *CHR* of *Sesbania rostrata* previously reported are ambiguous and should be re-identified; (c) IFSs share high levels of Sequence identity, indicating highly conserved during proteins evolution in plants; (d) *IFR*-*like* genes in *S*. *flavescens* and *Lupinus angustifolius* were clustered with homologs in nonlegume, sharing high sequence similarity and similar protein structures. These findings suggest the functional diversification of the multigene families, due to gene duplications in legume. Overall, our results offer reasonable gene annotations and comparative and functional analysis of legumes, as well as insights into the genetic evolution of the legume-specific biosynthetic genes of isoflavonoids.

## Methods

### Bioinformatics analysis

As no genome are available for *S*. *flavescens*, we constructed a *de novo* transcriptome assembly by using newly generated root RNA-seq data (unpublished). These *CHS*, *CHR*, *IFS* and *IFR* homologs in *S*. *flavescens* were identified by performing BLASTN from the BLAST+ package (Camacho *et al*., 2009) against our RNA-seq database. For each gene performance analysis, sequences of homolog genes reported in other legumes were used as queries and retrieved from NCBI database. For example, to identify *IFR* candidate genes of *S*. *flavescens*, *IFR* genes from *Medicago sativa* (accession number X58078.1) (Paiva *et al*., 1991) and *Pisum sativum* (accession number S72472.1) (Paiva *et al*., 1994) were blasted against our RNA-seq library.

Genome-wide analyses of CHS homologs in *M*. *truncatula*, *L*. *japonicus*, *G*. *max* and related nonlegumes have been identified (Zavala and Opazo, 2015). In our study, these sequences were re-conformed and located on the precise position of chromosome in legumes (https://bioinformatics.psb.ugent.be/plaza/versions/plaza_v3_dicots/). *CHR*, *IFS* and *IFR* homologs in *M*. *truncatula*, *L*. *japonicus*, *G*. *max*, *V*. *unguiculata*, *P*. *vulgaris*, *C*. *cajan* and *V*. *angularis* were mainly obtained from PlantGDB database (http://www.plantgdb.org/) and Legume information system (https://legumeinfo.org/). Phytozome, annotated with KOG, KEGG, ENZYME and pathways, was also included to attain broader coverage. In *Lupinus angustifolius*, homologs of CHR and IFS were blasted against recent released reference genome sequence (Hane *et al*., 2017). As reference analysis, related homologs of nonlegumes were retrieved from GenBank. All genes and accession numbers used were listed in Table 1.

**Table 1.**
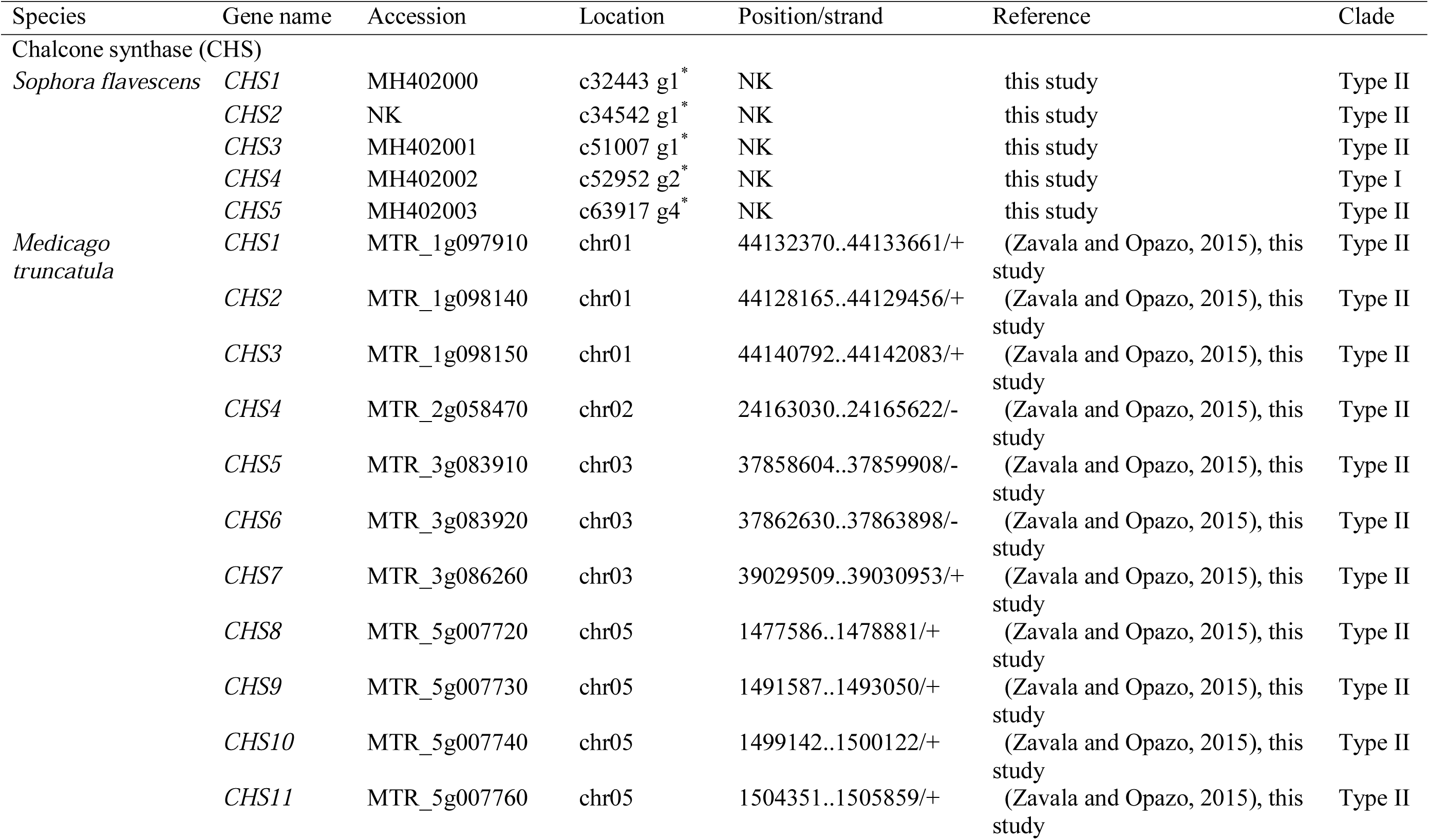
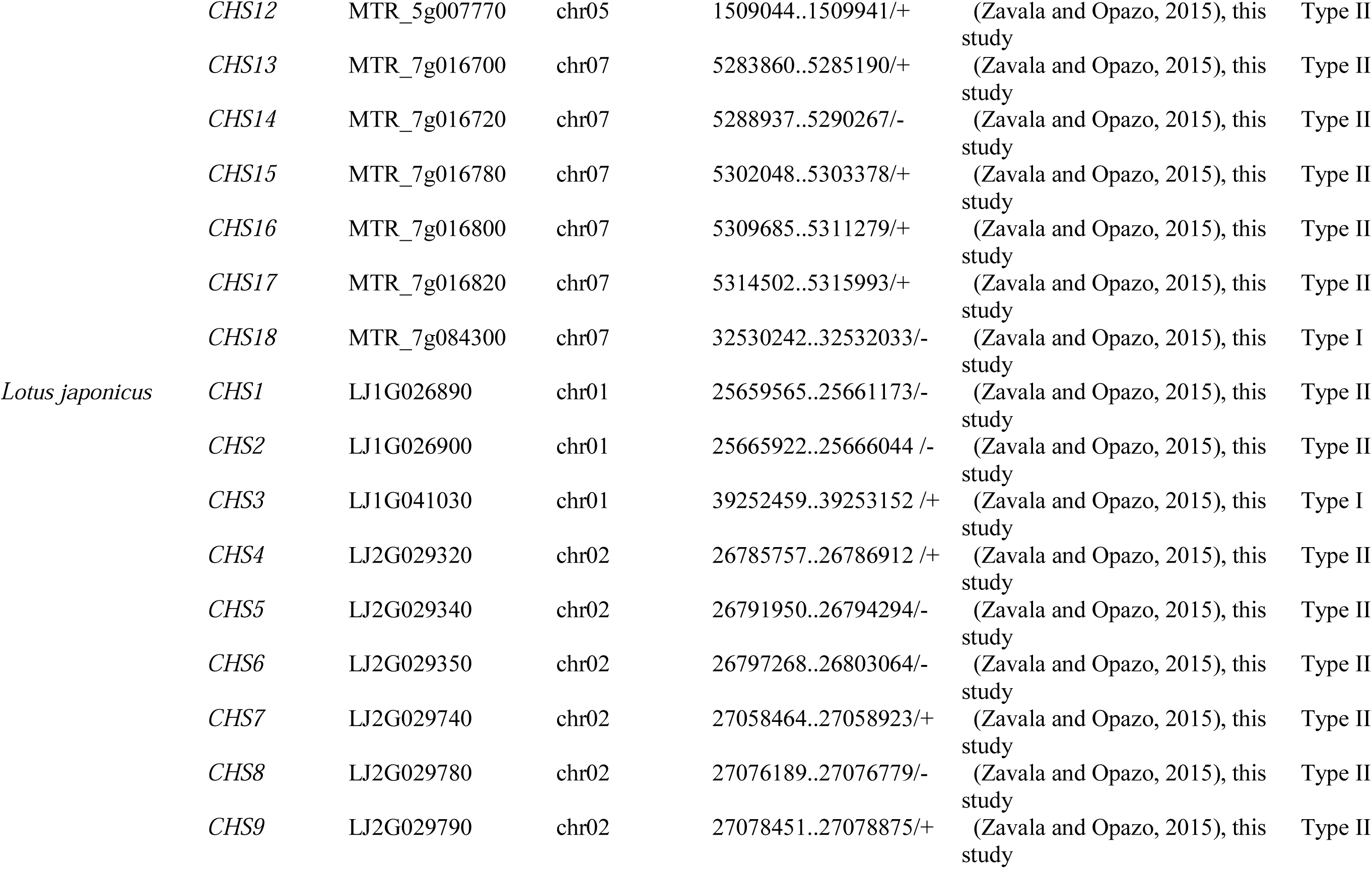
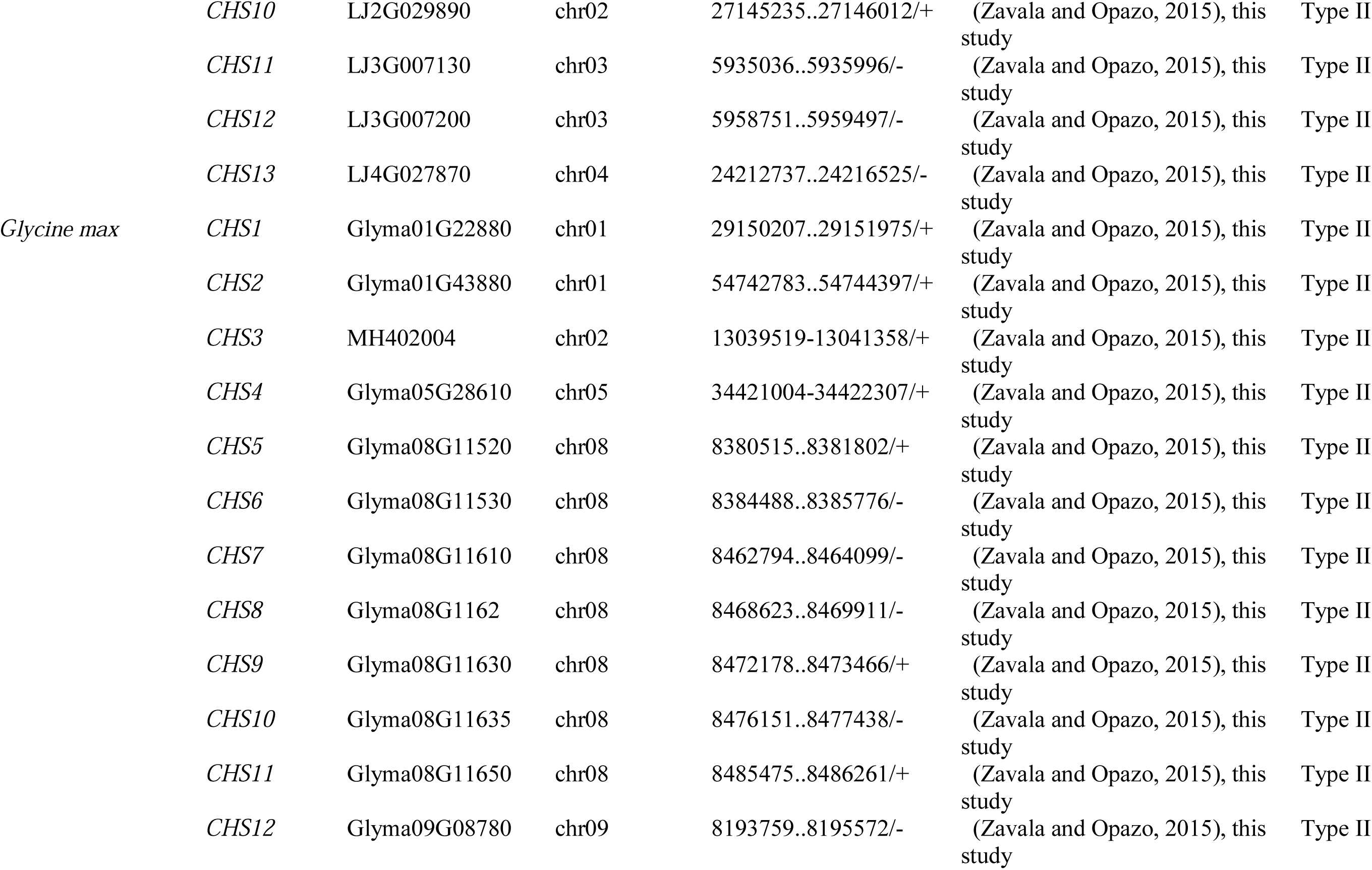
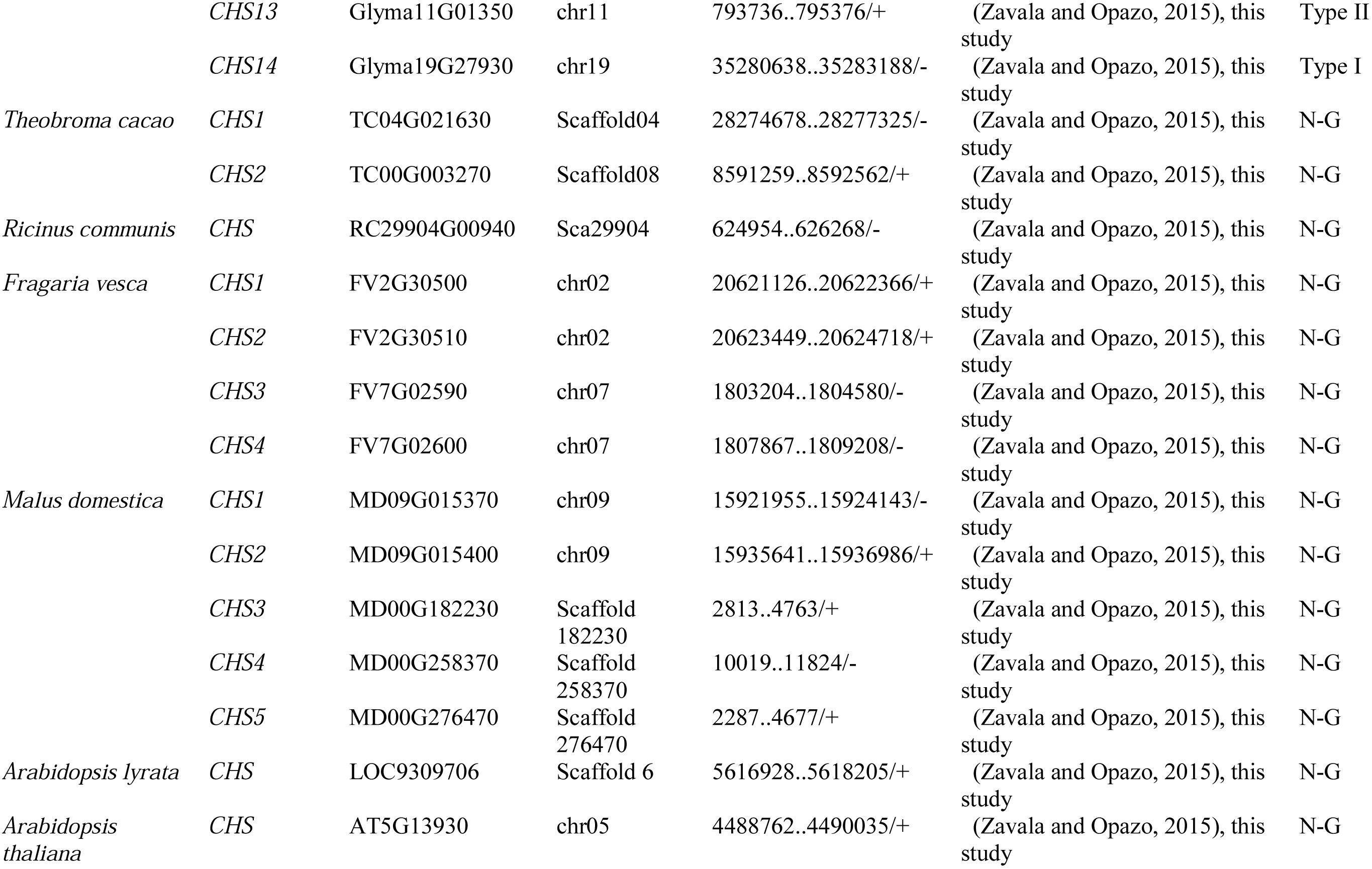
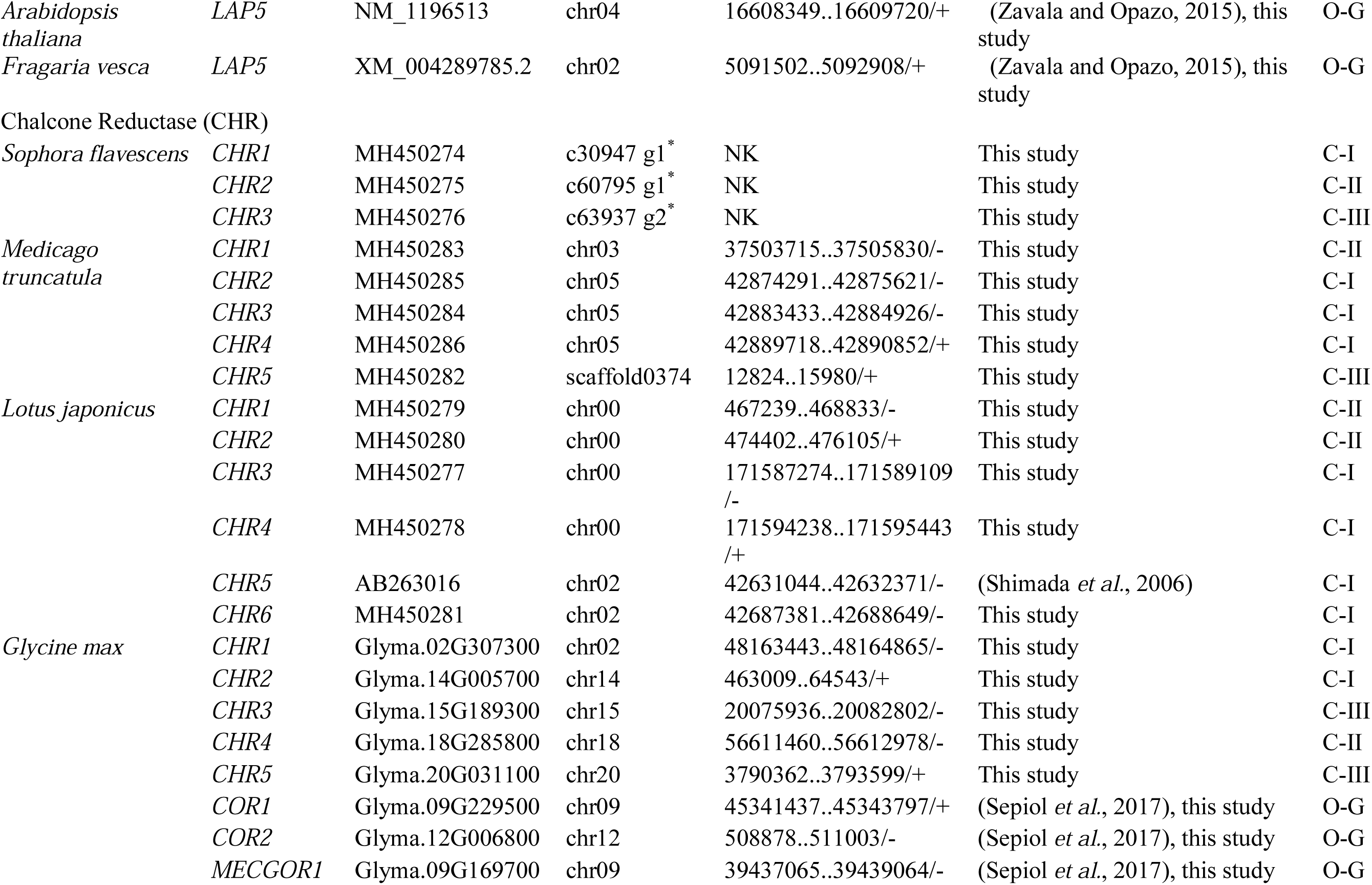
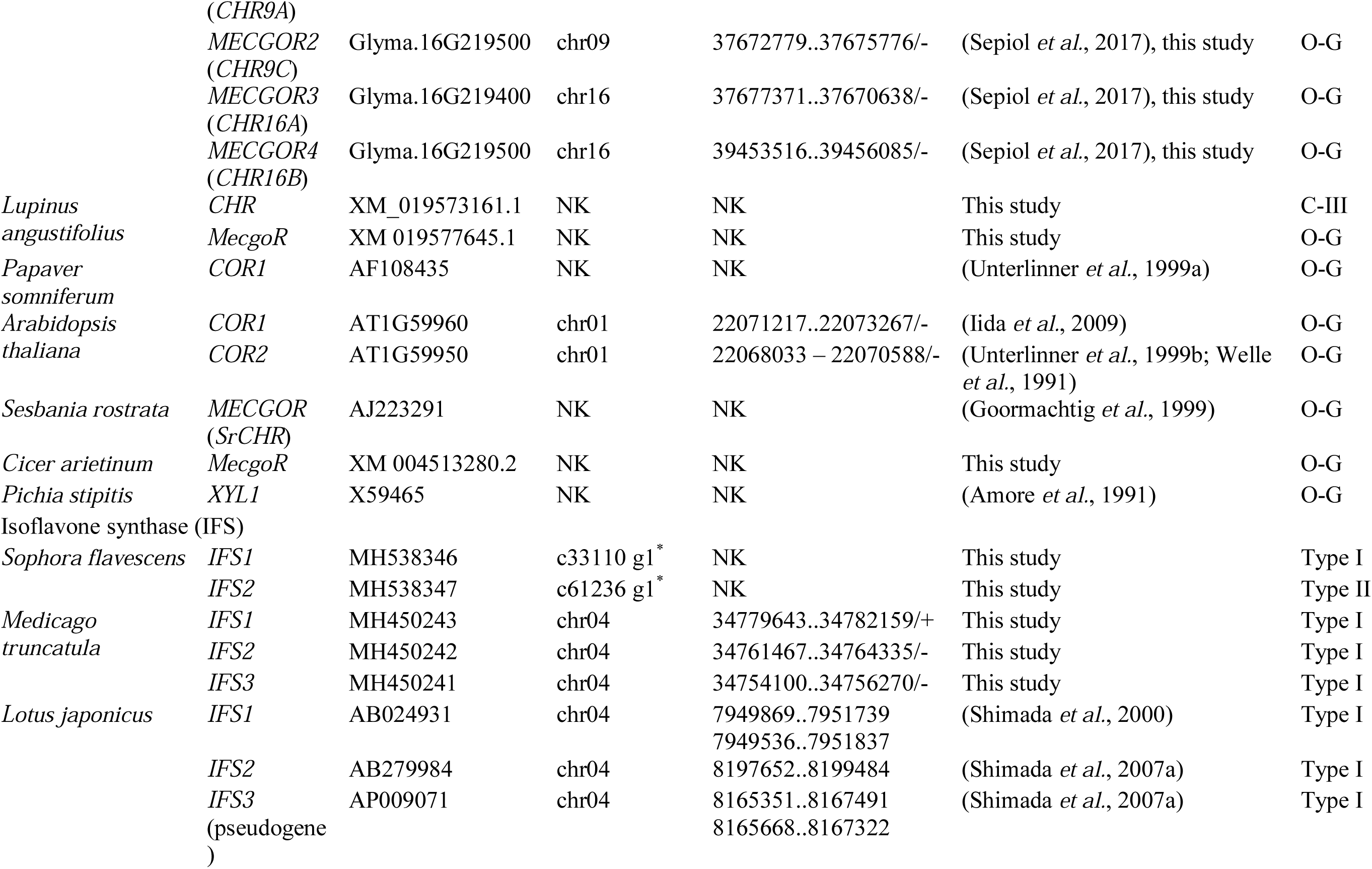
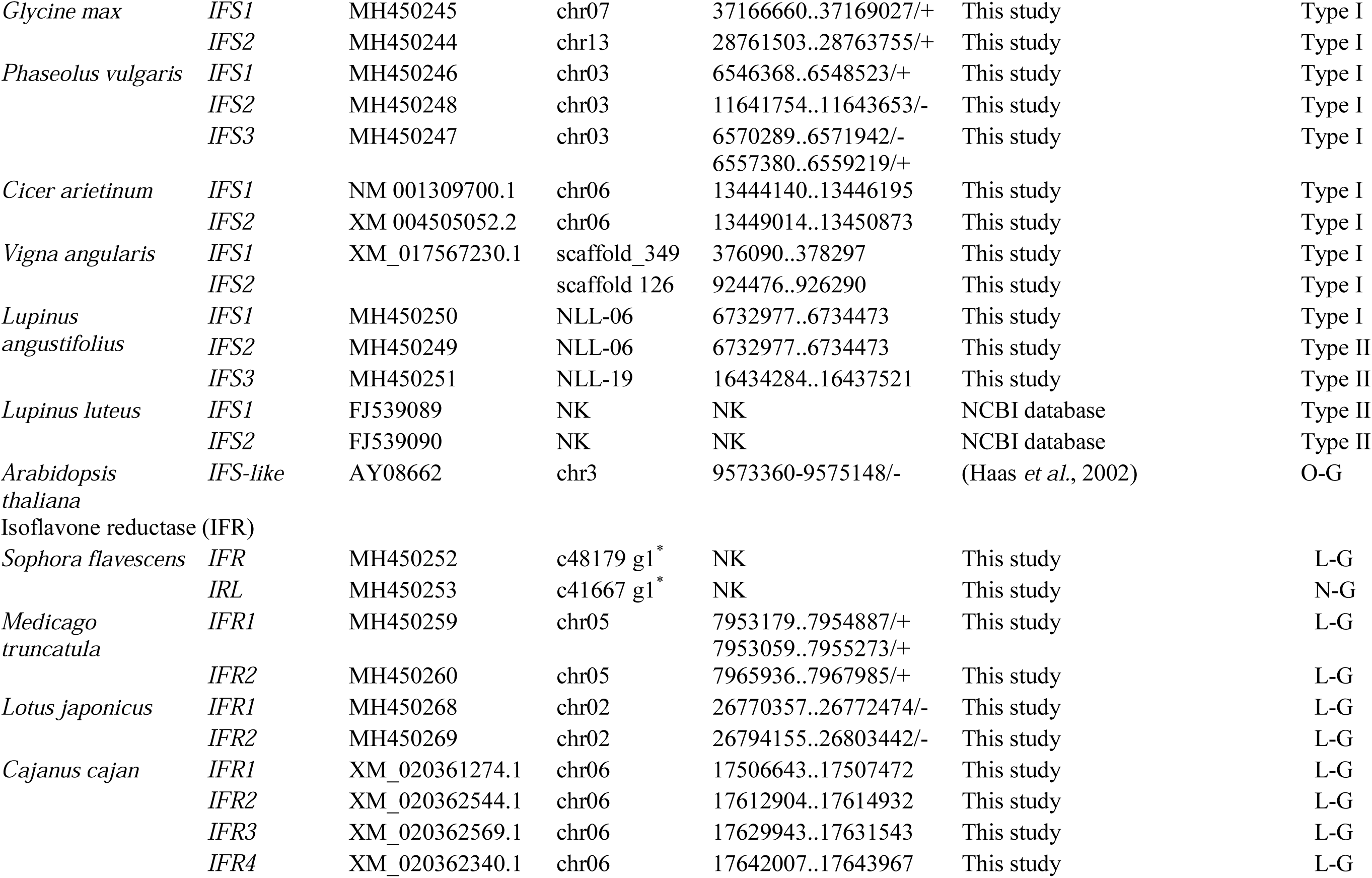
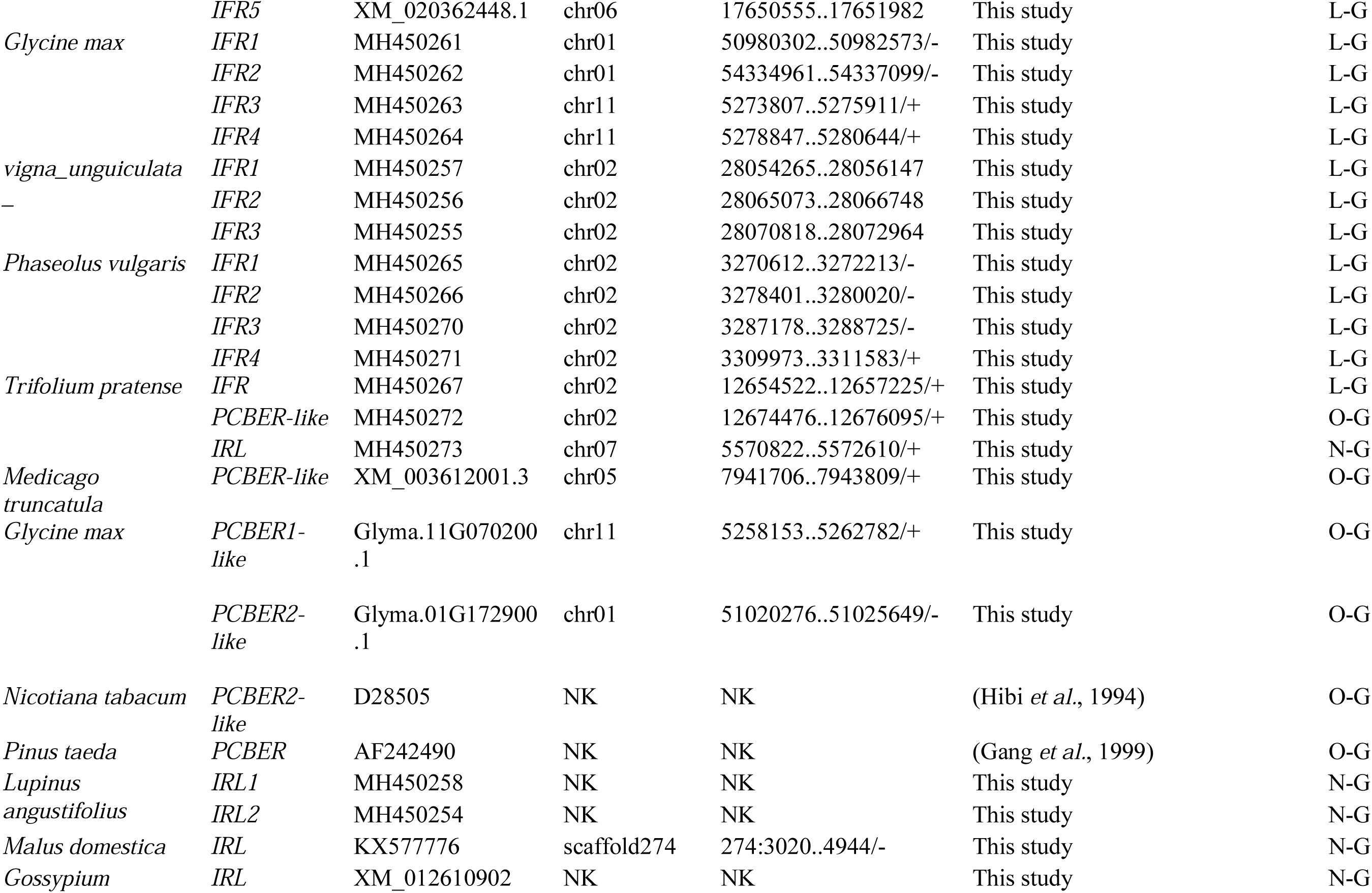
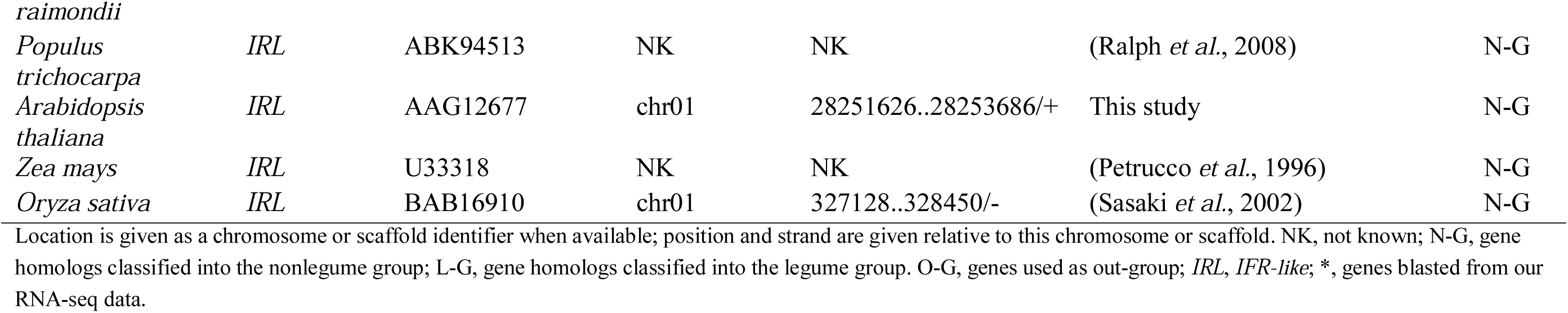
Summary of four key genes involved in (iso)flavonoid biosynthesis in legumes and related nonlegumes.

### Sequence similarity and phylogenetic analysis

Evolutionary similarity of *CHSs* nucleotide sequences between legume-specific and nonlegume groups was conducted in MEGA7 (Kumar *et al*., 2016). The estimates were obtained by a bootstrap procedure with 1000 replicates. Analyses were conducted using the Maximum Composite Likelihood model. All positions containing gaps and missing data were eliminated. The nucleotide sequences of *CHS*, *CHR*, *IFS* and *IFR* genes were aligned manually. The alignment lengths were 1173 bp for *CHSs*, 1077 bp for *CHR*, 1620 bp for *IFSs* and 1412 bp for *IFRs*, respectively. The evolutionary relationship of these four genes were inferred by using the Maximum Likelihood (ML) method based on the Tamura-Nei model (Tamura and Nei, 1993) integrated in MEGA 7. Initial trees for the heuristic search were obtained automatically by applying Neighbor-Joining and BioNJ algorithms to a matrix of pairwise distances estimated using the Maximum Composite Likelihood (MCL) approach, and then selecting the topology with superior log likelihood value.

### Comparative analysis of protein sequence

The protein sequences of IFS and IFR homologs in all studied species were translated from coding sequences. The alignment was conducted using ClustalX 2.1 and checked manually. As we did not obtain the whole coding sequence of IFS2 of *S*. *flavescens*, it was excluded in the further protein sequence alignment. The secondary structure elements in IFR structure were predicted according to the previous result (Wang *et al*., 2006).

## Results

### Identification of *CHS*, *CHR*, *IFS* and *IFR* homologs

We focus on four essential enzymes, chalcone synthase (CHS), chalcone reductase (CHR), isoflavone synthase (IFS) and isoflavone reductase (IFR), involved in the pathway of isoflavonoid biosynthesis in legumes. In order to shed more light on evolutionary route of four enzymes in legumes belonging to different phylogenetic Papilionoideae tribes, genomic resources of the inverted repeat loss clade (IRLC) of the Hologalegina (*Medicago truncatula*,), robinioids (*Lotus japonicus*), millettioids (*Glycine max*, *Phaseolus vulgaris* and *Vigna angularis*) and basal genistoids (*Lupinus angustifolius* and *Sophora flavescens*) were investigated in our study. As no genome data of *Sophora flavescens* is available, RNA sequencing (RNA-seq) and de novo assembly were performed to obtain the repertoire of isoflavonoids biosynthesis genes in *S*. *flavescens*. Total 137828 unigenes representing the total root transcriptome of *S*. *flavescens* were generated by RNA-seq (unpublished data). Meanwhile, homologs of *CHS*, *CHR* and *IFR* in some nonlegumes from public genomic data were also identified for comparative analysis.

### Multiple duplications of *CHS* homologs in legumes

Similar to *CHS* genes (*CHSs*) reported in other legumes (Zavala and Opazo, 2015), *CHSs* in *S*. *flavescens* also have many copies that *CHR1* (c32443 g1), *CHR2* (c34542 g1), *CHR3* (c51007 g1), *CHR4* (c52952 g2), *CHR5* (c63917 g4) were identified from root library (Table 1). To compare phylogenetic evolution of the *SfCHSs* with those in other legumes and nonlegumes, we download *CHSs* sequences that have been annotated (Zavala and Opazo, 2015) and further conformed the *CHSs* positions in genomes of different plant species by blasting (http://www.plantgdb.org/). The phylogenetic tree was reconstructed by using maximum likelihood (ML) method based on nucleotide sequences (Fig. 2). It showed that the copy number of putatively functional *CHSs* in legumes is higher than those in nonlegumes. For example, 18 *CHSs* and 1 *CHS* copy were present in *M*. *truncatula* and *A*. *thaliana*, respectively (Fig. 2). The result suggested that *CHSs* in legumes underwent more duplication events than those in nonlegumes. Meanwhile, most *CHSs* were phylogenetically grouped into a divergent legume-specific clade, named type II (Fig. 2). Intriguingly, *SfCHS4*, together with *CHS14* in *G*. *max*, *CHS3* in *L*. *japonicus* and *CHS18* in *M*. *truncatula*, was distributed to one of ancestral nonlegumious lineages (type I) that also contained two *CHS* homologs in *A*. *thaliana* and *A*. *lyrata*. Further, nucleotide sequences identities of representative *CHSs* in both legume-specific and nonlegume groups were calculated in every combination (Table S1). The high sequence identities (about 80%) within the legume-specific type II *CHSs* showed a close orthologous relationship each other. In contrast, the nucleotide identities between the two types of *CHSs* in legumes were significantly lower 75%. Taken together, these observations supported the hypothesis that the evolution of *CHSs* in legumes was derived from an ancestral repertoire presented in the last nonlegumious ancestor (Fig. 1).

**Fig.2.**
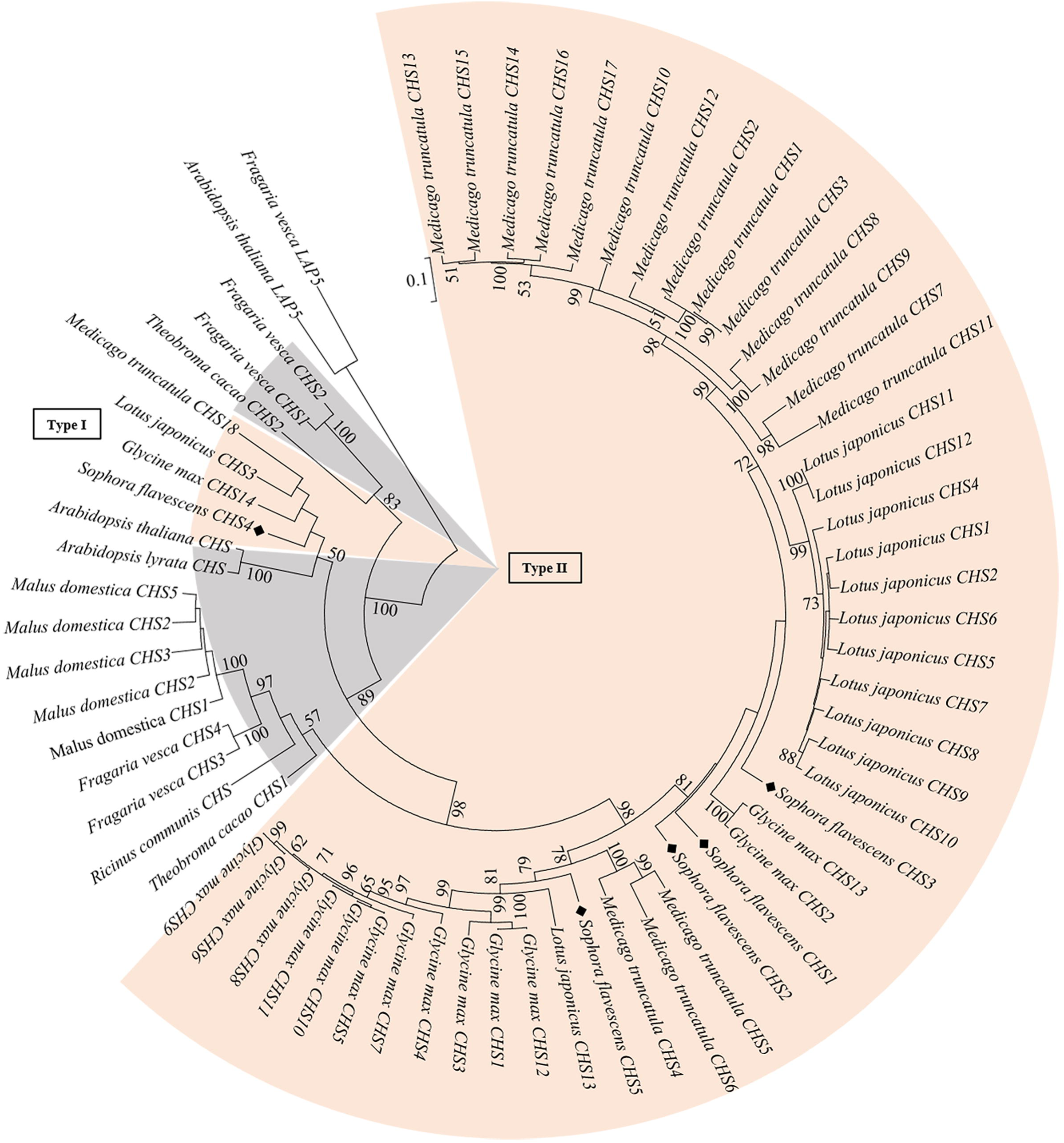
phylogenetic analysis suggested the origin of the legume-specific type II of *CHSs*. Maximum likelihood phylogenetic tree was constructed based on nucleotide sequences. Most of *CHSs* in legumes were grouped together, named type II. *CHS4* of *S*. *flavescens*, *CHS14* of *G*. *max*, *CHS3* of *L*. *japonicus* and *CHS18* of *M*. *truncatula* were classified into the type I group, closed with homologs in nonlegumes. Nude and gray colours indicate genes identified in legume and nonlegume, respectively. Black rhombus indicate *CHSs* of *S*. *flavescens*. *LAP5* of *F*. *vesca* and *A*. *thaliana* were used as an out-group. Only bootstrap values greater than 50% are shown at branch nodes. Bar, 10% nucleotide substitution per site.

### *CHR* homologs in legumes as AKR4

Three putative *CHR* genes (*CHRs*) in *S*. *flavescens*, named *SfCHR1*-*SfCHR3*, were identified using BLAST against our RNA-seq library. To figure out copy number variations and phylogenetic relationship of *CHRs* among different legume species, *CHR* homologs in *L*. *japonicus*, *M*. *truncatula* and *G*. *max* were searched from Legume Genome Database (https://legumeinfo.org/). Besides, other related genes such as genes encoding codeinone reductases (COR) were selected as reference. These nucleotide sequences in all studied species were aligned for building phylogenetic tree (Fig. 3). The result showed that *CHRs* in legumes were grouped together in a distinguished subclade of ARK4 family, indicating highly conserved in the evolutional history. *COR* homologs in nonlegumes were out-grouped from clade of *CHR*, constituting another subclade of ARK4 family. Further analysis showed *CHRs* from these four legume species fell into 3 distant subclades (Fig. 3). In clade I, *MtCHR2*-*MtCHR4*, located on chromosome 5, shared high similarity at nucleic acid lever (Table 1). Besides *SfCHR1*, *GmCHR1*-*GmCHR2* and *LjCHR3*-*LjCHR6* were gathered together, respectively. Some *CHR* homologs such as *SfCHR2* and *MtCHR1* were placed into divergent subclades belonging to Clade II. *LjCHR1*, distributed in clade II, showed catalytic activity in a previous study (Shimada *et al*., 2007a), giving evidence that other *CHRs* in this clade may also encode active enzymes. Notably, clade III as an early branch of *CHRs* was firstly investigated. These data suggested that *CHRs* in legumes were possibly duplicated twice before the speciation of Papilionoideae in the evolutionary processes.

**Fig.3.**
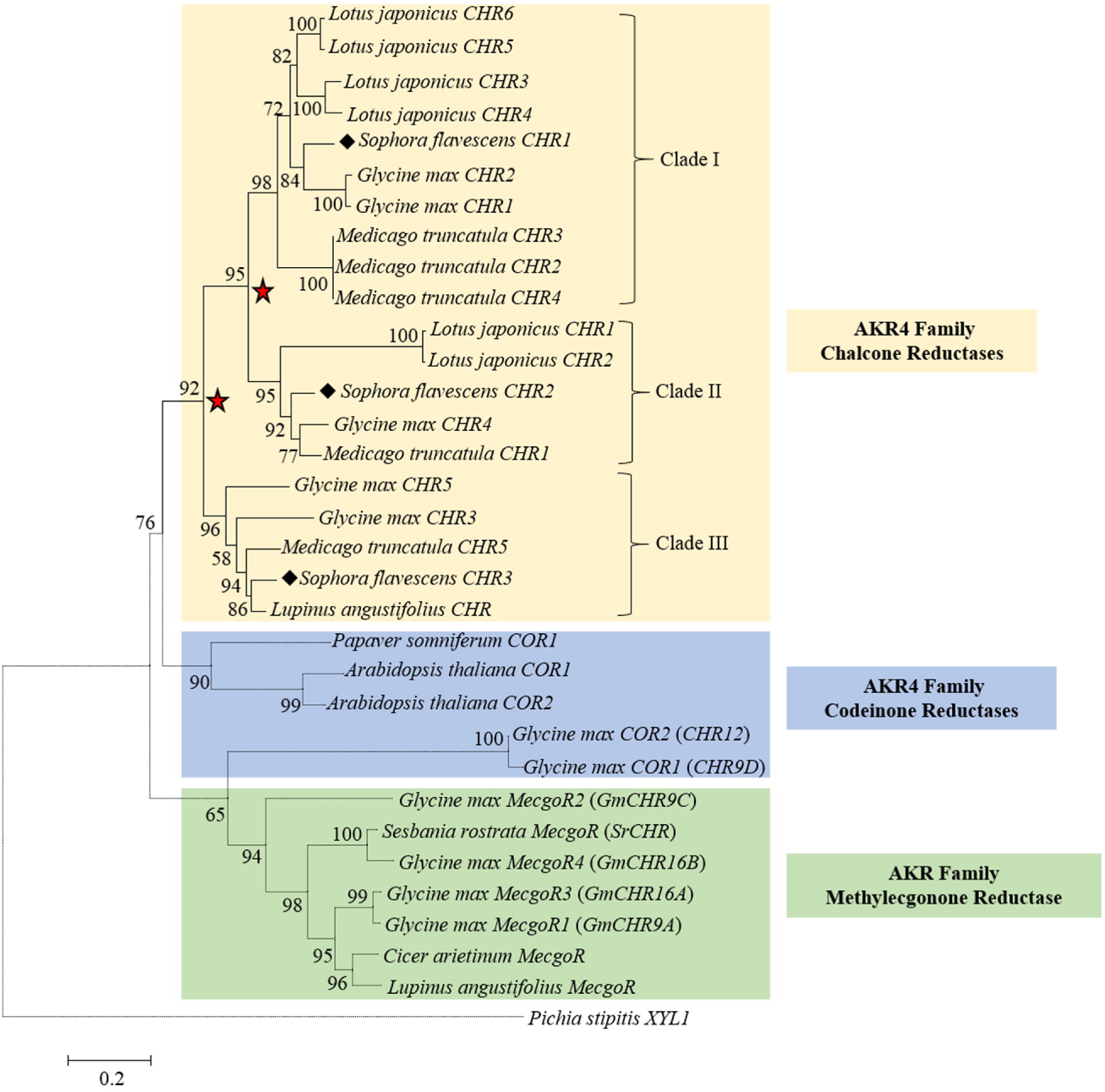
*CHRs* in legumes belonging to AKR4 Family were grouped together. The phylogenetic analysis showed that these *CHRs* in legumes were distributed into three subclades, clade I, II and III, suggesting that *CHRs* were possibly duplicated twice (indicated by five-pointed star) before the speciation of Papilionoideae in the evolutional processes. Some *CHR* homologs in *G*. *max* and *CHR* of *S*. *rostrata* were re-identified based on phylogenetic evolution. Three colours from up to down indicate chalcone reductases, codeinone reductases and methylecgonone reductase of AKR4 subfamily members, respectively. Dotted branch indicates phylogenetic clades except for *CHRs*. Black rhombus indicate *CHRs* of *S*. *flavescens*. XYL1 of *P*. *stipitis* was used as an out-group. Only bootstrap values greater than 50% are shown at branch nodes. Bar, 20% nucleotide substitution per site.

Recently, *CHRs* in *G*. *max* were investigated and have 11 genes copy, which distributed in multiple clades of AKR family (Sepiol et al., 2017). However, *CHR9A*, *CHR9C*, *CHR9D*, *CHR12*, *CHR16A* and *CHR16B* from *G*. *max* were distantly placed out of the *CHRs* clade in our study. To further conform the identities of these 6 *CHRs*, we blasted them in the NCBI database. We found that *CHR9D* and *CHR12* were annotated as codeinone reductase–like protein and shared more than 85 % nucleotide sequence similarity with known *COR* homologs in legumes. More supersizing, *CHR9A*, *CHR9C*, *CHR16A* and *CHR16B* have highest similarity with genes coding methylecgonone reductase (MecgoR) in *C*. *arietinum* and *L*. *angustifolius*. Furthermore, similar case was also observed in *S*. *rostrata* (Goormachtig et al., 1999) that *SrCHR* gene (AJ223291.1) was clustered into *MecgoR* group (Fig. 3). Based on our blasting results and phylogenetic analysis, this gene was supposed to be a *MecgoR*-*like* gene. Overall, our study provides strong evidences for identifies of *CHRs* in AKR4 family and helps us to distinguish phylogenetic relationship between *CHRs* and other closed genes like *MecgoRs* and *CORs* in legumes.

### Conserved evolution of IFS homologs in legumes

Isoflavone synthase (IFS), also named as 2-HIS, introduces the isoflavonoid biosynthetic pathway in legumes (Wang, 2011). Compared with many copies of *SfCHR* and *SfCHS*, only two *IFS* candidate genes, *SfIFS1* and *SfIFS2*, were identified from the same root RNA-seq library. The blast analysis in NCBI database showed that *SfIFS1* and *SfIFS2* have 80% and 72.6% at nucleotide level with well-studied *GmIFS1* and *GmIFS2*, respectively. We further identified *IFS* homologs (*IFSs*) from genomes of *P*. *vulgaris*, *V*. *angularis*, *G*. *max*, *M*. *truncatula*, *C*. *arietinum* and *L*. *japonicus* (Table 1). A *IFS* phylogenetic tree was then constructed and showed that most homologous of legumes, named type I, were grouped together, forming a main phylogenetic clade (Fig. 4). Meanwhile, five *IFS* candidate genes including *SfIFS2* and four *IFSs* in *L*. *angustifolius* and *L*. *luteus*, named type II, were distantly grouped in an early-branching clade. To further figure out the evolutional relatedness of two types of IFSs, all encoded IFS protein sequences were aligned and showed a high degree of similarity (Fig. 5), ranging from 70.0% to 100.0%, indicating that the evolution route of IFSs in legume was distinctly conserved. This observation was also supported by motif analysis that all studied IFSs shared conserved “I” helix, FSAGTDST, proposed to be involved in oxygen binding by P450 proteins. In addition, the heme-iron ligand signature, FGSGRRMCPG, were also exhibited at the C-terminal sequences of IFSs (Steele et al., 1999). We also found two obvious sequences distinctions among these proteins; one particular region was that two high conserved residues in type II of IFSs (20 P and 55 L) was lost in corresponding region of type I; another one showed high variation of amino acid sequences from position 423 to 435 of type I but highly conserved in type II. Further studies will be needed to investigate the functional difference of two types of IFSs.

**Fig.4.**
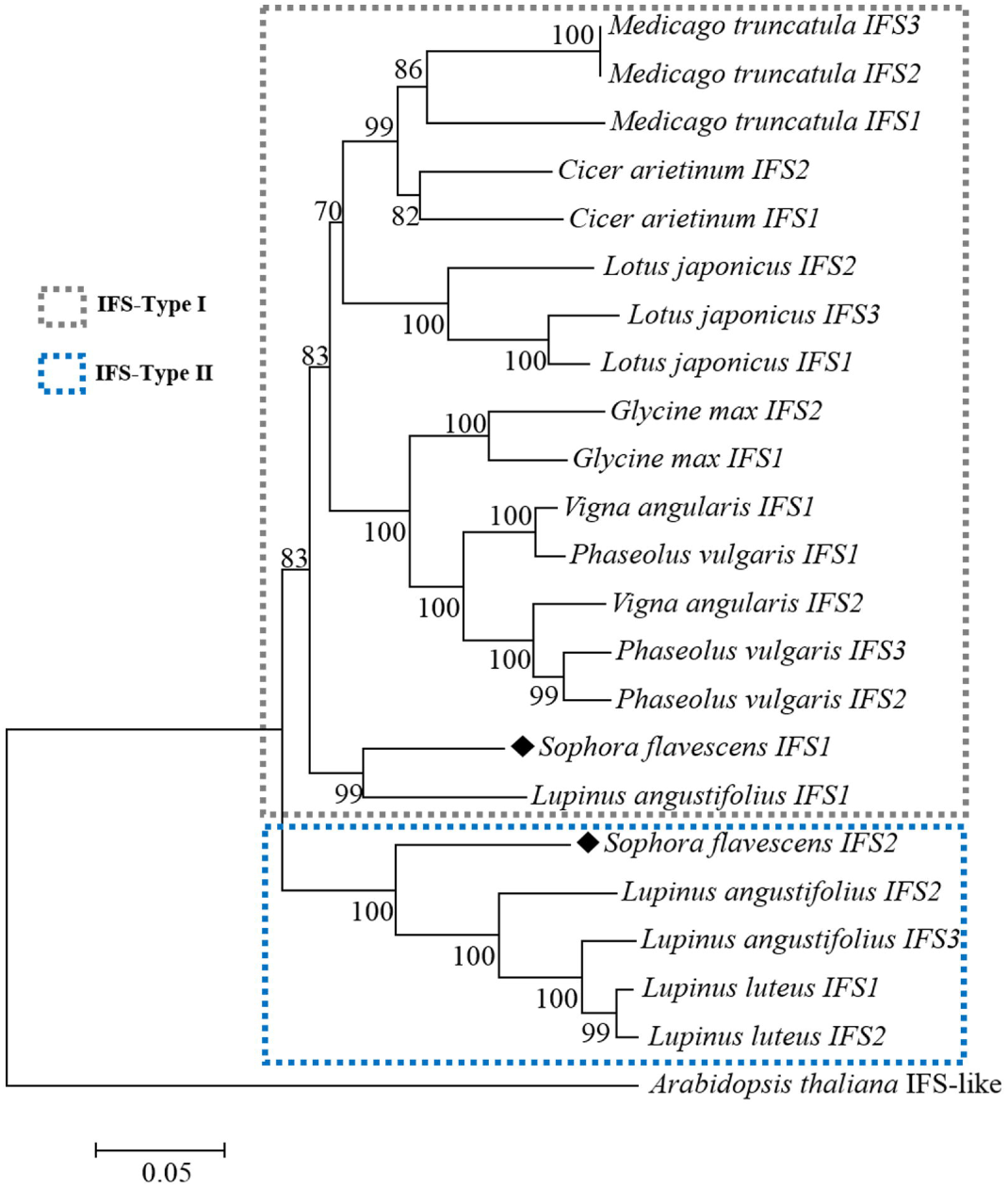
Maximum likelihood phylogenetic tree of IFSs in legume. Most IFS homologs of legumes, named type I (grey dotted box), were grouped together, forming a main phylogenetic clade. In addition, five IFS candidate genes in *S*. *flavescens*, *L*. *angustifolius* and *L*. *luteus*, were distantly grouped in an early-branching clade, named type II (green dotted box). Black rhombus indicate IFSs of *S*. *flavescens*. IFS-like gene of *A*. *thaliana* was used as an out-group. Only bootstrap values greater than 50% are shown at branch nodes. Bar, 5% nucleotide substitution per site.

**Fig.5.**
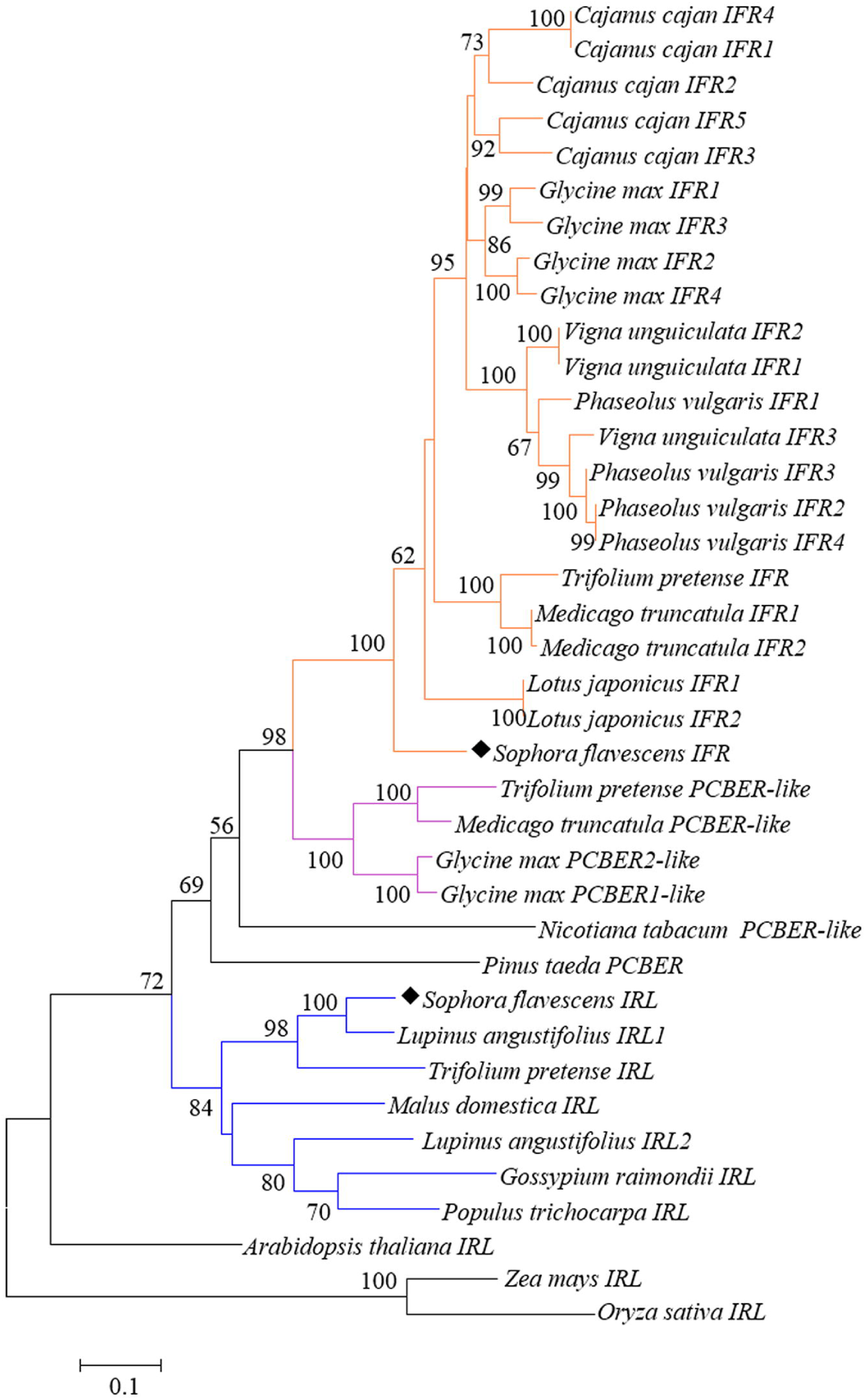
Phylogenetic analysis of *IFR* homologs in legumes and nonlegumes. The deduced nucleotide sequences of putative *IFRs* were aligned and the Maximum Likelihood (ML) tree was constructed using MEGA7 software. The orange branch indicates phylogenetic position of most legumious *IFRs*. *PCBER* candidate genes (in purple branch) in legumes may be evolved from the same ancient clade with IFRs. *IFR*-*like* (*IRL*) genes of *L*. *angustifolius*, *T*. *pretense* and *S*. *flavescens* were clustered together with IFR homologs in nonlegumes such as *IFR*-*like* gene in *G*. *raimondii* (in blue branch). Black rhombus indicate IFRs of *S*. *flavescens*. *IFS*-*like* genes in *A*. *thaliana*, *Z*. *mays* and *O*. *sativa* were used as an out-group. Only bootstrap values greater than 50% are shown at branch nodes. Bar, 10% nucleotide substitution per site.

### Evolutional analysis and characterization of protein sequences between IFR and IFR-Iike (IRL) in legumes

The obtained cDNA clone *SfIFR* and *SfIRL* were 1539 and 1693 nucleotides in length, encoding proteins of 309 and 320 amino acids, respectively. The analysis of conserved protein domains (http://www.ncbi.nlm.nih.gov/Structure/cdd/wrpsb.cgi) showed that *SfIFR* contained an N-terminus characteristic of NAD(P)-binding domain and a small C-terminal domain presumed to be involved in substrate binding. On the other hand, SfIRL belonged to NmrA-like family and displayed a Rossmann-fold NAD(P)H/NAD(P)^+^ binding (NADB) domain, which was consistent with the observed protein structure of IFR in *M*. *sativa* (Wang et al., 2006).

To gain insight into the phylogenetic diversity of *IFR* genes (*IFRs*) in legumes and their related genes in nonlegumes, database searches were firstly conducted. A considerable number of genes sharing high nucleotide sequence identity with *SfIFR* and *SfIRL* were identified, many of which were annotated as encoding isoflavone reductase-like proteins. ML phylogenetic tree indicated that *SfIFR* grouped with most leguminous *IFRs* together, which constituted a monophyletic legume-group (Fig. 6). we further observed *PCBER*-*like* genes from *G*. *max*, *M*. *truncatula* and *Trifolium pretense* were grouped together, forming a closed sister clade with the legume-specific clade. Interestingly, *SfIRL*, *IRLs* in *L*. *angustifolius* and *T*. *pretense* were evolved from same early-branch with *IRLs* in nonlegumes such as *Gossypium raimondii*. In order to investigate potential divergence between these four unusual *IRLs* in legumes and *IFRs* in legume-specific clade, amino acid sequences alignment of representative IFRs, PCBERs and IRLs were performed. The secondary structure elements predicted in the IFR structure were shown in Fig. 7. We observed a conserved G17-X-X-G20-X-X-G23 fingerprint region presented in the N terminus (loop β1-α1) of all IFRs and related proteins. The largest difference between IFRs in Legume-group and IFRs in nonlegume-group was observed in the region between strand β2 and helix α2. Compared with SfIFR and other leguminous homologs, SfIFL, LaIRL1 and LaIRL2 have ten less amino acid residues deletion in this region as well as in IRLs in nonlegumes (Fig. 7).

**Fig.6.**
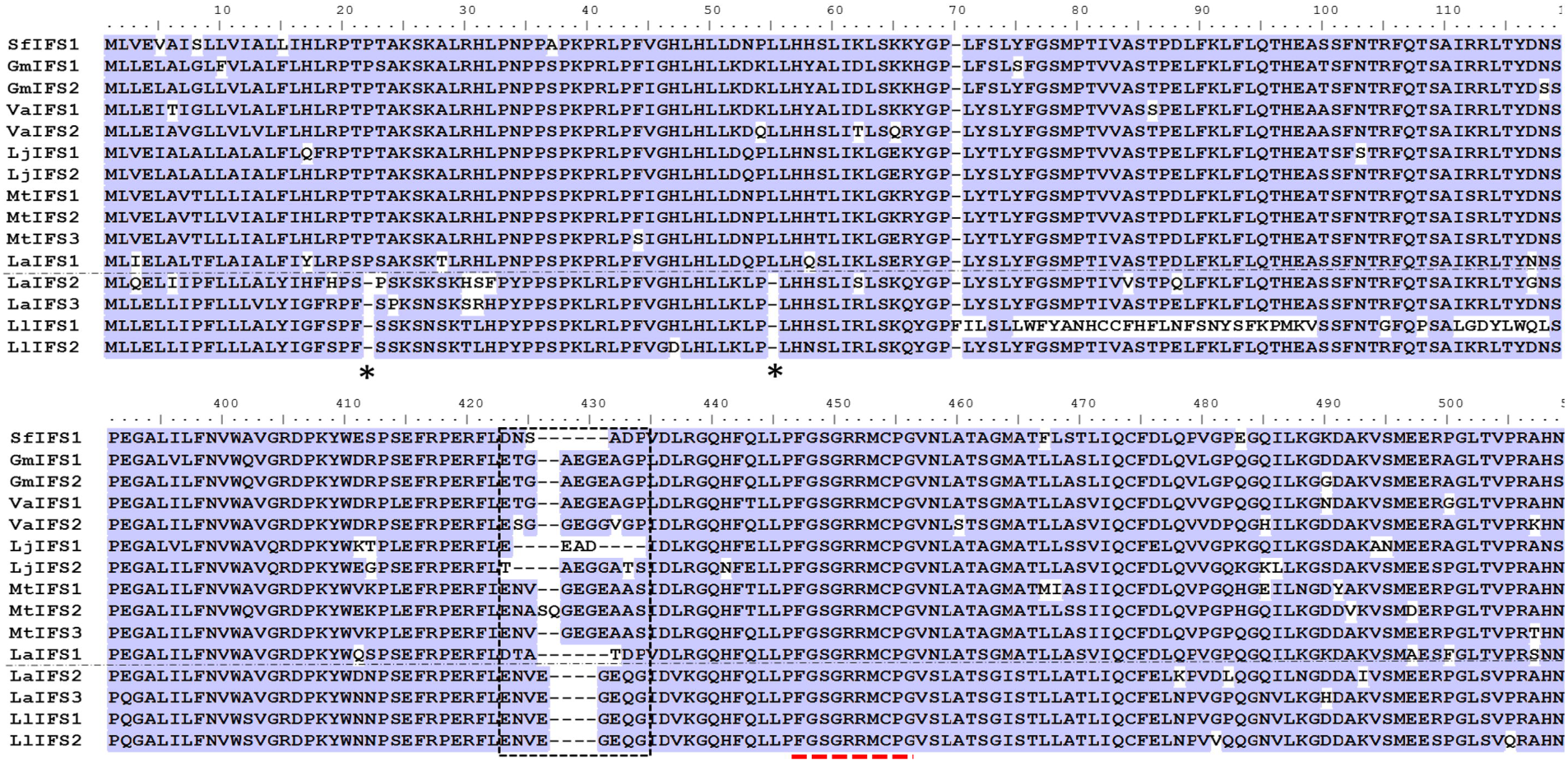
Sequences alignment of IFSs. Protein Sequences were from type I group (above the dotted line) including *S*. *flavescens* (SfIFS1), *G*. *max* (GmIFS1 and GmIFS2), *V*. *angularis* (VaIFS1 and VaIFS2), *L*. *japonicus* (LjIFS1 and LjIFS2), *M*. *truncatula* (MtIFS1 and MtIFS2) and *L*. *angustifolius* (LaIFS1); type II group (below the dotted line) including *L*. *angustifolius* (LaIFS2 and LaIFS3) and *L*. *luteus* (LlIFS1 and LlIFS2). Identical residues are highlighted. ^∗^, two conserved residues lost in type II of IFSs. Black dotted box indicates the drastic variation of amino acid sequences in IFSs. Red dotted line indicates the heme-iron ligand signature of IFSs.

**Fig.7.**
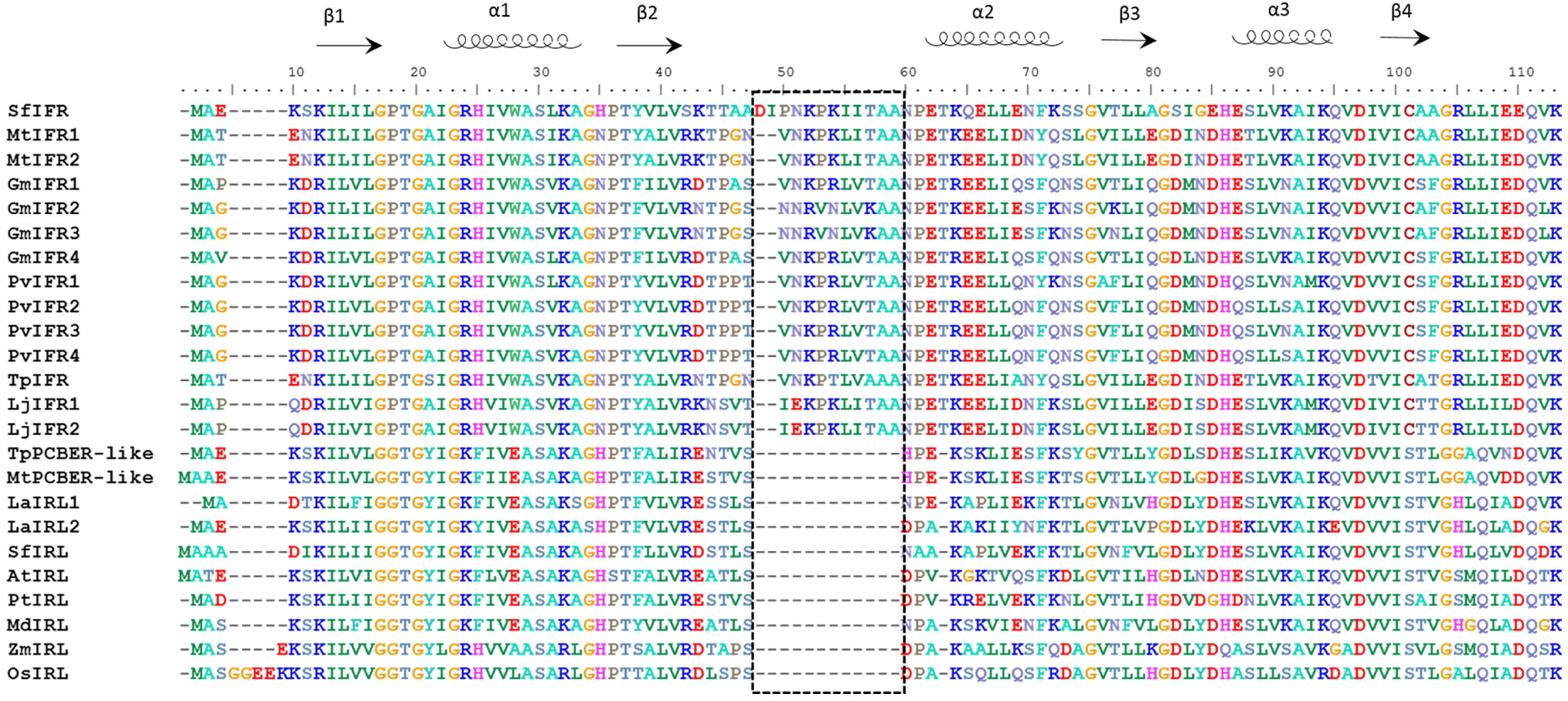
A notable distinction between IFRs and related IFR-like homologs in legumes and nonleugmes was observed in the region between strand β2 and helix α2 (black dotted box). Sequence alignment of IFRs from *S*. *flavescens* (SfIFR), *M*. *truncatula* (MtIFR1 and MtIFR2), *G*. *max* (GmIFR1, GmIFR2, GmIFR3 and GmIFR4), *P*. *vulgaris* (PvIFR1, PvIFR2, PvIFR3 and PvIFR4), *T*. *pratense* (TpIFR), *L*. *japonicus* (LjIFR1 and LjIFR2); PCBER-like proteins in *T*. *pratense* (TpPCBER-like) and *M*. *truncatula* (MtPCBER-like); IFR-like (IRL) proteins from *L*. *albus* (LaIRL1 and LaIRL2), *S*. *flavescens* (SfIRL), *A*. *thaliana* (AtIRL), *P*. *trichocarpa* (PtIRL), *M*. *domestica* (MdIRL), *Z*. *mays* (ZmIRL) and *O*. *sativa* (OsIRL). The secondary structure elements observed in the IFR structure are shown above the alignment.

## Discussion

We investigated the evolutionary history of key enzymes involved in the legume-specific isoflavonoid biosynthesis in *Sophora flavescens* and other representative species that located in divergent phylogenetic lineages of Papilionoideae, a subfamily of Fabaceae. Phylogenetic analysis revealed that *CHSs* and *CHRs* composed multigene subfamilies and formed divergent gene subclades in many cases. *IFSs*, only found in legumes, were distinctly conserved in the evolutional history. We also found that some *CHS* and *IFR* paralogous copies in legume existed as non-leguminous type. Based on these results, we propose that gene duplication events and neofunctionalization of some key enzymes in the isoflavonoid biosynthesis is a major driving force for the evolutionary gain of a legume-specific function such as inducer of Rhizobia nodulation. This hypothesis can be convinced by the observation that two rounds of duplication in the MtLYK3-LjNFR1a clade are ancestral to evolutionary gain of a legume-specific nod factor receptor (De Mita et al., 2014).

The phylogeny of chalcone synthase gene (CHS) in *Rosids* has been studied (Shimada *et al*., 2007b; Zavala and Opazo, 2015). However, none of these studies was specifically focused on the evolutional origin of legume-type of *CHS*. Instead, they only considered multiple duplication processes occurred in the legume-specific clade of *CHSs*. In *L*. *japonicas*, *CHS1* was phylogenetically distant with other *CHS* homologs in legumes (Shimada et al., 2007b). Here, we found the same case also existed in *S*. *flavescens*, *G*. *max*, and *M*. *truncatula* (Fig. 2). These finding suggested that legume-specific *CHSs* might be derived from the ancestral *CHS* genes in nonlegume-clade. Additional evidence supporting this hypothesis has emerged from the study about two types of chalcone isomerase (*CHI*) genes, involved in the biosynthesis of general flavonoids and legume-specific 5-deoxy(iso)flavonoids in *L*. *japonicus* (Shimada et al., 2003). Therefore, it is likely that type I CHS in nonlegumes may only function in the production of 6-deoxychalcone, whereas legume-specific type II CHS may involve production of both 6-deoxychalcone and 6-hydroxychalcone.

CHRs, belong to the aldo–ketoreductase (AKR) superfamily, play key roles in the biosynthesis of deoxychalcone-derived isoflavonoids such as soybean phytoalexins. *CHRs* belonging to three subclades in legumes were identified in our study, which was inconsistent with results that showed that only two subclades were observed (Sepiol et al., 2017). It was supposed that two independent duplication events occurred in the evolution process of *CHRs* in Papilionoideae (Fig. 3). The ancestral *CHR* in clade III was possibly duplicated once before the speciation of clade I and II, and the subsequent duplication event caused the paralogous genes in the two subclades. In our study, some *CHR*-*like* genes in *G*. *max* and *S*. *rostrata* were re-identified as *COR*-*like* and *MecgoR*-*like* genes, showing that *CHRs*, *CORs* and *MecgoRs* were very close to each other. Further study is required to illustrate the biochemistry and functional divergence of CHRs, CORs and MecgoRs, which will provided more evidences for gene classification.

Isoflavone synthase (IFS) is the key enzyme for biosynthesis of isoflavones in legumes. By introducing soybean IFS1 into *A*. *thaliana* which had no ability to synthesize isoflavone, genistein accumulated at a different level (Liu et al. 2002 and 2000). Based on our results, the duplication event of *IFS* genes in legume was conserved. Exceptionally, we also found that three and two *IFS* homologs identified in *L*. *angustifolius* and *S*. *flavescens* were classified into type I and II groups, which placed in distant phylogenetic clades. P450s and NADPH-P450 reductases are dissociated and independently anchored on the outer face of the endoplasmic reticulum (ER) by amino-terminal hydrophobic anchors (Werck-Reichhart and Feyereisen, 2000). At the N-terminal sequence, targeting sequence such as LLELAIGLVVLALFLHLR (GmIFS1) was supposed to be required for proper orientation in the ER membrane (Yamazaki *et al*., 1993). In our study, conservation analysis revealed that amino acid 20P of IFSs, near ER region, was highly conserved in Type IFS-I while was lost in type *IFS*-*II*. Further investigation is required to shed more light on evolutional mechanism and enzymatic activity of two IFS versions in legumes.

The most intriguing and unique feature of *IFR* homologs was that some *IFRs* present in *S*. *flavescens*, *L*. *angustifolius* and *T*. *pretense* had close relationship with *IRLs* in nonlegumes. *SfIRL*, *TpIRL*, *LaIRL1 and LaIRL2* were classified into nonlegume group based on phylogenetic analysis as well as amino acid alignment. We also observed the missing residues at the 5’ region of IFRs, PCBERs and IRLs in all studied plants. This observation was consistent with the result that IFRs were comparable with PCBERs and were deduced to have the same overall basic structure, a continuous NADPH-binding domain and a smaller substrate-binding domain (Min *et al*., 2003). The highly structure similarity suggested that SfIRL and LaIRLs may play similar function with IRLs in nonlegumes. However, the function information of these IRL proteins present in non-legume plants are not available. Because both IFR homologs of *L*. *angustifolius* were IRL type and we did not find any legume-specific homologs in *L*. *angustifolius* from available genome and RNA-seq data. This raises the question of whether this key step of isoflavonoid biosynthetic pathway in *L*. *angustifolius* is different from that in other legume species. However, our results can not rule out the functional possibility of isoenzymes. Further studies about biochemical function difference of two types IFR homologs in *S*. *flavescens* are needed. Overall, our results suggest that evolution of legume specific *IFRs* and homologs in nonlegumes may be forced from common ancestral gene repertoire. Considering the fact that the dominated role of isoflavonoids in the process of symbiosis between rhizobia and legumes, more efforts are needed to investigate isoflavone components secreted by *L*. *angustifolius* and other plants when interaction with rhizobium or pathogens.

## Acknowledgements

This work was financially supported by the National Natural Science Foundation of China (31770039). Y. J. and Y. Z. acknowledge support from China Scholarship Council. The authors declare that they have no conflict of interest.

## Author Contribution

Y. J. designed the research, and collected and analysed the data. Y. Z. assisted in developing the research design. Y. J wrote the manuscript and W. C. edited the manuscript.

